# Deep Lineage: Single-Cell Lineage Tracing and Fate Inference Using Deep Learning

**DOI:** 10.1101/2024.04.25.591126

**Authors:** Mehrshad Sadria, Allen Zhang, Gary D. Bader

## Abstract

Recent advances in single-cell RNA-sequencing and lineage tracing techniques have provided valuable insights into the temporal changes in gene expression during development, tumour progression, and disease onset. However, there are few computational methods available to analyze this information to help understand multicellular dynamics. We introduce Deep Lineage, a novel deep-learning method for analyzing time-series single-cell RNA-sequencing with matched lineage-tracing data. Our method accurately predicts early cell fate biases and gene expression profiles at different time points within a clone, surpassing current state-of-the-art methods in fate prediction accuracy. Additionally, through in silico perturbations in cellular reprogramming and hematopoiesis development data, we show that Deep Lineage can accurately model dynamic multicellular responses while identifying key genes and pathways associated with cell fate determination.

## Introduction

Developmental biology aims to understand the processes by which single cells divide and differentiate into diverse cell types in different tissues (1). Single-cell RNA-sequencing (scRNA-seq) is a powerful tool for investigating developmental landscapes by profiling gene expression in individual cells at different stages (2). While scRNA-seq is valuable for investigating genetic mechanisms in cell differentiation, its destructive nature restricts it to capturing a single time point snapshots of a developing system. This constraint poses challenges in tracing lineage relationships and tracking cell trajectories over time (3).

To overcome this limitation, a range of methods have been developed to study temporal relationships between cells. One class of computational methods, trajectory inference, predicts the order of cells along a developmental path based on the idea that cells that are developmentally related will exhibit similar gene expression profiles (4). However, trajectory inference methods are limited in their capacity to capture the complexity of developmental processes, such as lineage branching and cellular plasticity, and can be negatively affected by technical factors and noisy data (5,6). New experimental lineage tracing technologies that use CRISPR/Cas9 genome editing to record cell division events provide another avenue for studying cell development (7). Using data produced by these technologies, computational methods have been developed to infer developmental trajectories by considering both cell lineage and cell state information from multiple time points (8–11). Integration of these two types of information has made predictions more accurate (12). However, despite these advances, the existing methods still can not accurately predict cell fate (13).

Here, we present Deep Lineage, a novel computational model that integrates lineage tracing and scRNA-seq data to learn cellular trajectories, enabling the prediction of gene expression and cell type at any given time point along a trajectory. Deep Lineage uses Long Short-Term Memory (LSTM), Bi-directional Long Short-Term Memory (Bi-LSTM) or Gated Recurrent Units (GRUs) to model complex sequential dependencies and temporal dynamics of a cellular trajectory. An autoencoder-learned embedding captures essential features of the data to simplify input to the LSTM. To assess method performance, we analyzed data sets containing both lineage tracing and scRNA-seq data. Our results demonstrate that Deep Lineage surpasses state-of-the-art methods in detecting early fate bias within a developmental process and can accurately predict gene expression profiles at held-out time points within a clone during a biological process. Furthermore, we show Deep Lineage’s ability to identify important genes and pathways involved in cell fate decision-making during a developmental process and accurately simulate genetic perturbation effects on gene expression of cell progenies. Such insights can aid in designing experiments to reprogram cells into specific cell types and potentially translate to clinical applications.

## Results

### Model Overview

Deep Lineage uses lineage tracing and multi-timepoint scRNA-seq data to learn a robust model of a cellular trajectory such that gene expression and cell type information at different time points within that trajectory can be predicted. It can be used to predict single-cell time points that have not been measured in a clone, including predicting cell fate at future time points from early time points (early cell fate bias). An autoencoder-learned embedding captures essential data features and a LSTM, Bi-LSTM (14) or GRU (15) is used to support cell type classification and gene expression regression tasks (Fig. 1). Deep Lineage treats cells and their progenies within a clone as interconnected entities. Drawing inspiration from natural language processing (NLP) (16), we conceptualize cellular relationships in terms of “clones” which represent cells ordered within a shared lineage (like words in a sentence) and gene expression relationships (like words that have similar meaning) (13).

**Fig. 1:**
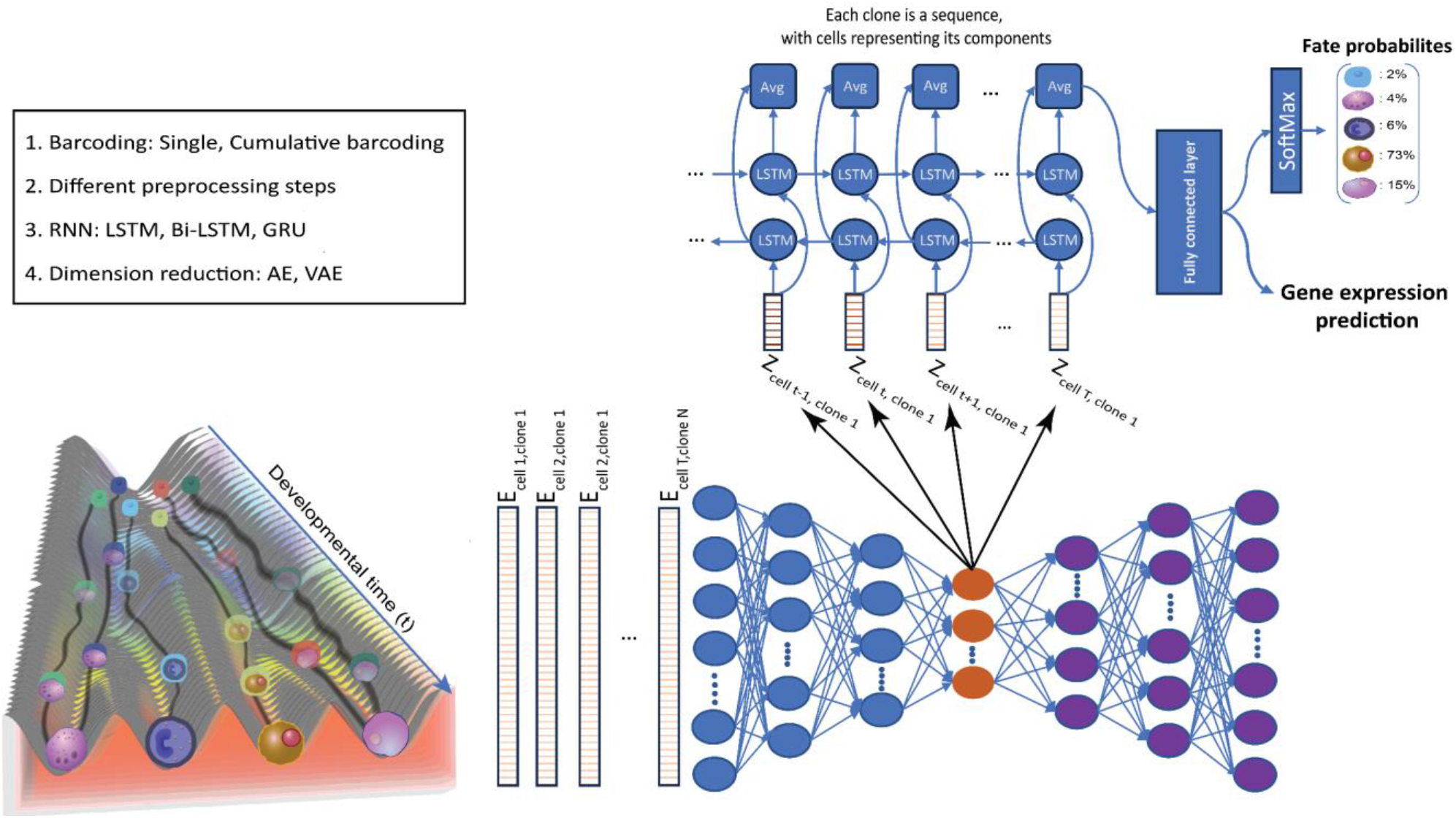
Predicting Gene Expression and Early Cell Fate Bias via Combined scRNA-seq and Lineage Tracing. A visual depiction of the Deep Lineage is presented. On the left, the Waddington landscape illustrates the developmental trajectory, showing how cells within each clone (represented by different colours) differentiate into distinct mature cell types originating from stem cells. On the right, by combining scRNA-seq data and lineage tracing information the gene expression profiles of cells within each clone are used as input for the autoencoder. The resulting latent embeddings of cells in a clone are subsequently used as inputs for the LSTM, Bi-LSTM, or the GRU model. This integrated architecture enables the accurate prediction of both early cell fate bias and gene expression profile of unseen days within a clone. Notably, Deep Lineage offers the flexibility to adapt different preprocessing steps, diverse barcoding techniques, various dimensionality reduction methods, and a range of various deep learning models (left black box).

We train the model, including an extensive hyperparameter search and data preprocessing to optimize the model’s performance (lowest loss for reconstruction and classification), on predicting held-out time points. We then train the optimized model with all data. To use the model, single-cell gene expression profiles from one or more time points are encoded in the learned autoencoder’s low-dimensional representation. This encoding and the cell’s clone label are fed into the LSTM or GRU network to predict cell fate bias for input cells, by examining cell types for each clone at a predicted future time point (Fig. 1). The model also predicts gene expression profiles for cells at different stages of the trajectory that can be used to impute missing time points. Deep Lineage can incorporate lineage tracing data using either single barcoding or cumulative barcoding methods (Supplementary Figure 1). Lineage tracing with cumulative barcoding significantly boosts the accuracy of the model compared to single barcoding resulting in more reliable predictions for both early cell fate bias and gene expression patterns at different stages of development (see below). Then by applying the SHapley Additive exPlanations (SHAP) method (17) to the trained model, Deep Lineage can accurately identify key genes and regulatory mechanisms linked to different fate outcomes at different developmental stages.

### Deep Lineage accurately predicts gene expression and early cell fate bias in mouse hematopoiesis

We applied Deep Lineage to the published lineage tracing data of mouse hematopoiesis, which consists of gene expression of 130,887 cells, and 25,289 genes (18). These clones are sampled from three time points, on days 2, 4, and 6. We focused on the main cell lineage in this data set, specifically neutrophils, monocytes, and their progenitors (Fig. 2a). We trained models to predict the gene expression profile of held-out cells at two-time points within a clone (predict day 4 by training with days 2 and 6 and predict day 6 by training with days 2 and 4)(Fig. 2b), including optimizing preprocessing steps and hyperparameters (see methods). We calculated the correlation between the gene expression profile of the predicted and held-out cells. The results demonstrated a high degree of similarity (*R* = 0.81 for both experiments), indicating that Deep Lineage accurately predicts the gene expression of cells unseen by the model on both day 4 (trained on days 2 and 6) and day 6 (trained on days 2 and 4) (Fig. 2c and Supplementary Figure 3a). We also observe a strong positive correlation between the average predicted expression of all genes and actual expression in both monocytes and neutrophils on day 4 (Fig. 2d,e). Deep Lineage excels at predicting average gene expression and the full distribution of diverse genes, even for cells within a clone that were previously unseen by the model (Fig. 2f,g).

**Fig. 2:**
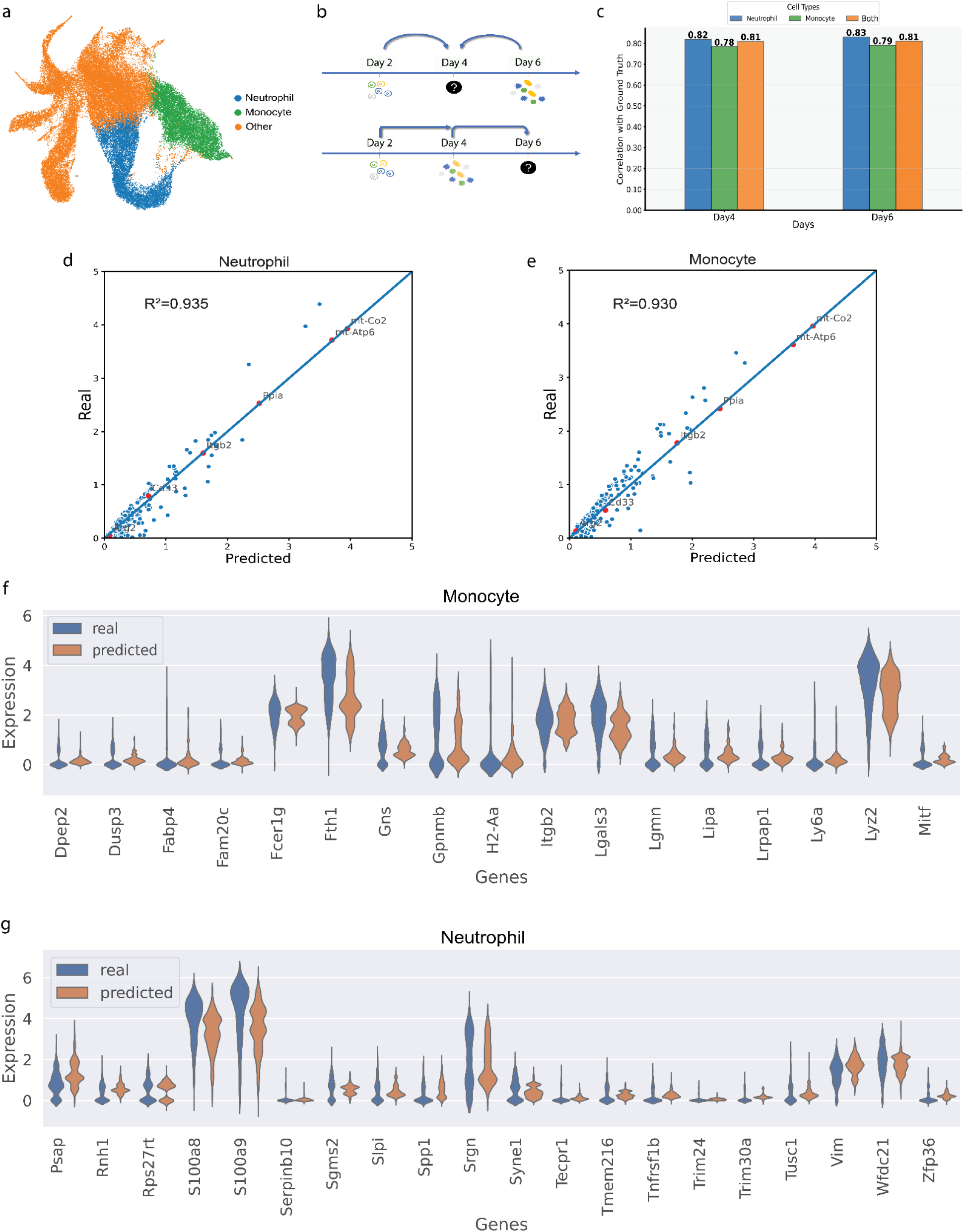
Deep Lineage accurately predicts single-cell gene expression of unseen cells in Hematopoiesis. a, UMAP plot of the hematopoiesis data from Weinreb et al. Each point represents an individual cell, color-coded by cell type. b, Schematic representation of the gene expression prediction process using two regression models: one for predicting day 4 gene expression (trained on data from days 2 and 6), and the other for predicting day 6 gene expression (trained on data from days 2 and 4) within a clone. c, Correlation of gene expression between Deep Lineage predictions and cells excluded during training, categorized by cell types and time points. d,e, Comparison of average gene expression values of 2000 genes between Deep Lineage predicted and real cells for Monocytes and Neutrophils. Gene expression at day 4 was averaged across predicted cells per clone, and then correlation was computed with actual data for each gene (*R*^*2*^ indicates the squared Pearson correlation coefficient between the predicted and ground truth values). f,g, Violin plots show gene expression distributions between predicted and real cells for randomly selected genes both Monocytes and Neutrophils.

To investigate early cell fate bias (Fig. 3a) in hematopoiesis, we trained a new bidirectional LSTM model using the cell type classification task and used it to predict late time point data from an early time point. Using just the gene expression of cells on day 2 of a clone, with the masking of data from days 4 and 6, the model attains a predictive accuracy of 76.8% in determining cell fate on day 6 (Fig. 3b). Providing gene expression of cells on days 2 and 4 of a clone improves the performance slightly and the model predicts the clonal fate on day 6 with 78.5% accuracy. If the model has access to the gene expression data from cells of all three days within a clone, it can almost perfectly predict the fate of the clone (99.2%) (Fig. 3b). Subsequently, we generate the area under the curve (AUC) for cell fate prediction of this model by using gene expression of cells on days 2 and 4, and all 3 days (Fig. 3c). Thus, by using the trained classifier, we can predict the early progenitor bias from an early time point, accurately differentiating between monocyte and neutrophil outcomes. The model performance improves when trained with additional time points. Furthermore, to investigate the importance of lineage tracing and clonal information, we conducted an ablation study involving random sampling of cells for each time point while disregarding the clonal information, for the train on day 2, predict day 6 experiment. The results indicate that neglecting lineage tracing information substantially reduces Deep Lineage’s prediction accuracy from 77% to 52%. Adjusting the model’s architecture (LSTM vs. GRU, hyperparameter changes) results in minimal change in accuracy (from 50% to 52%), highlighting the important role of clonal structure in fate prediction (Supplementary Figure 5a and Supplementary Table 3).

**Fig. 3:**
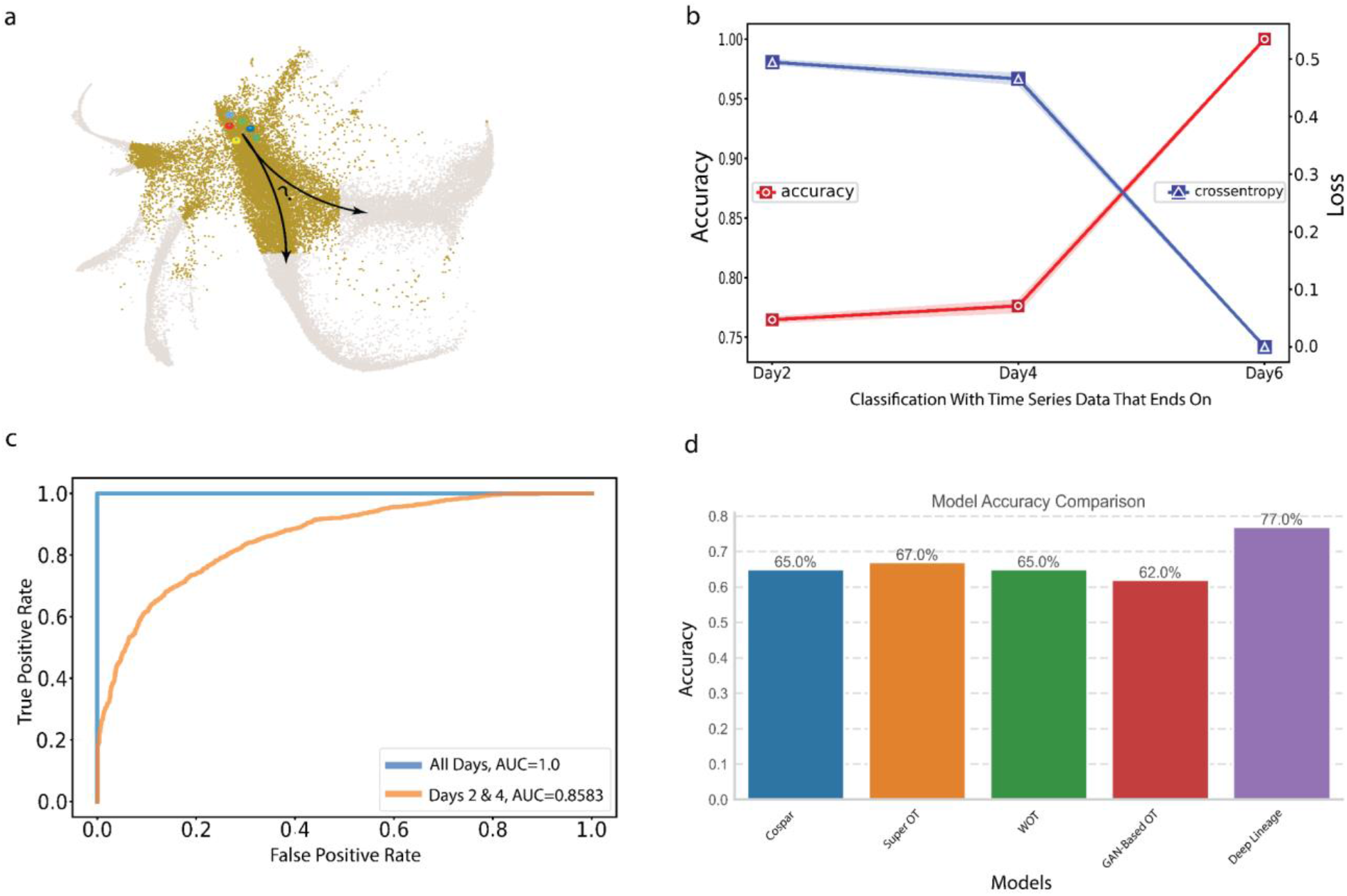
Exploring Progenitor Bias Prediction in Hematopoiesis with Deep Lineage and Comparison to State-of-the-Art Methods. a, Schematic UMAP diagram illustrating the trajectory of stem cell differentiation and the possible cellular outcomes with cell colors indicating distinct clonal lineages. Grey points represent other mature cells types (not neutrophils and monocytes). b, Performance evaluation of the classifier using accuracy and cross-entropy loss metrics to predict early cell fate bias by employing gene expression of cells on just day 2, using cells on days 2 and 4, and including all three days within a clone. c, Receiver Operating Characteristic (ROC) curves and Area Under the Curve (AUC) values for the model using cells from days 2 and 4 or all three days with a clone, showing our model’s performance in early cell fate bias prediction. Day 2 performance is very close to day 2&4 performance, thus not plotted. d, Comparative analysis with state-of-the-art methods in predicting early cell fate bias. Deep Lineage outperforms existing methods showing a higher classification accuracy.

Next, in a thorough comparison, we assessed our method alongside the state-of-the-art models: Cospar (9), Super OT (10), WOT (8), and GAN-Based OT (10), in predicting the fate of held-out clones during training. Our results indicate that Deep Lineage outperforms all other existing methods, demonstrating superior accuracy in predicting early fate bias (Fig. 3d). Specifically, the comparative analysis revealed that Deep Lineage achieved a high accuracy rate of 77%. In contrast, other methods such as Cospar attained a rate of 65%, Super OT exhibited a slightly higher accuracy at 67%, WOT maintained a similar accuracy of 65%, and GAN-Based OT achieved 62%.

### Deep Lineage accurately predicts gene expression and early cell fate bias in regeneration

To evaluate the performance of our model on another dataset with more time points, we applied Deep Lineage to the reprogramming process of fibroblasts into induced endodermal progenitor (iEP) cells over 28 days (19). This data comprises gene expression data for 18803 cells and 28001 genes, across six time points (days 6, 9, 12, 15, 21, 28), with corresponding barcode-derived lineage data at three time points (days 0, 3, 13) (Fig. 4a). Following the same approach as above, we first perform an extensive hyperparameter search to select the parameters that yield the highest correlation between the held-out samples during training and those predicted by the model (See methods, Supplementary table 2). Then we assess the ability of our model to predict gene expression on different days during the fibroblast transition to iEP cells. Our method consistently has high performance in generating realistic gene expression profiles, regardless of the specific time point chosen for analysis (Fig. 4b, Supplementary Figure 3b). We also compared the gene expression distribution between the model’s prediction and held-out cells at day 28 and showed that Deep Lineage can accurately predict the full gene expression distribution for both failed and reprogrammed cells (Fig 4c, and Supplementary Figure 4). Also, we observe a high similarity (*R*^*2*^ = 0.987) between the average predicted gene expression and the actual expression levels for genes in both successfully and failed reprogrammed cells (Fig. 4d,e).

**Fig. 4:**
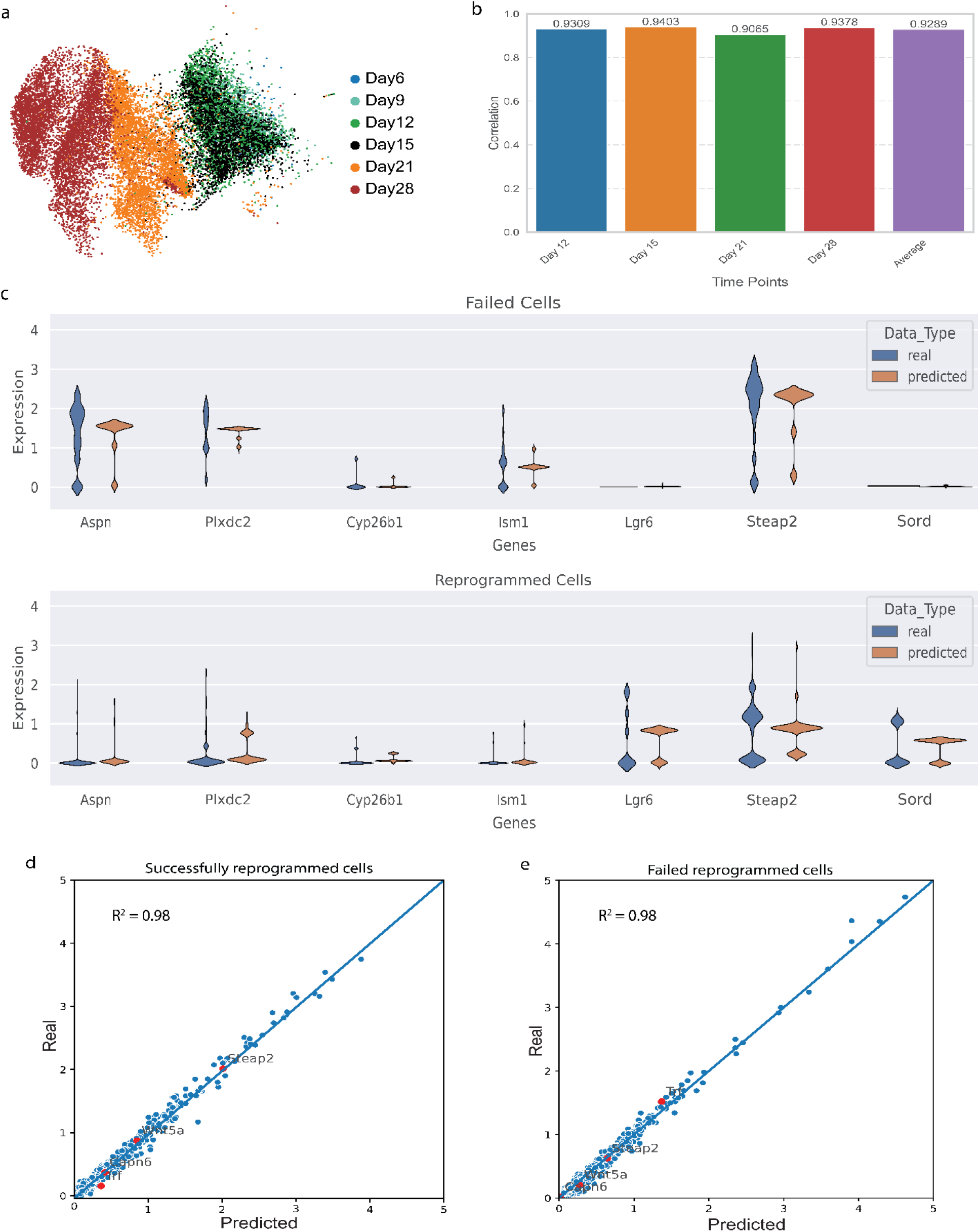
a, UMAP visualization demonstrates fibroblast cell reprogramming into iEPs, dots represent cells and are color-coded by time point. b, Correlation values between gene expression of ground truth cells and Deep Lineage predictions for cells excluded during training across different stages of the reprogramming process. c, Violin plots show gene expression distributions between real cells and Deep Lineage predictions for randomly selected genes for both Failed and Reprogrammed cells. d,e. Comparative analysis of the top 2000 differentially expressed genes between predicted and real cells. High correlation values are observed for both successful and failed reprogramming cells, highlighting the accuracy of the model for different cell types (*R*^2^ indicates the squared Pearson correlation coefficient between the predicted and ground truth values).

We evaluate Deep Lineage’s performance in predicting the outcome of the reprogramming process with ground truth labels and compare it to the current state-of-the-art method, Cellrank. (20). After training by providing the first four time points of the process (day 6, day 9, day 12, day 15) our model can predict the reprogramming outcome on day 28 with high accuracy (94%) (Fig. 5b). ROC analysis shows that Deep Lineage outperforms Cellrank irrespective of the number of days analyzed (Fig. 5c). As expected, by giving the model more early time points to use to infer later time points, the performance of both methods improved, but Deep Lineage performs better in all cases (Fig. 5c). To test our model under ablation conditions, we predict the reprogramming outcome by randomly sampling data points from days 6, 9, 12, 15, and 21 without considering clonal information. As seen for the mouse hematopoiesis ablation study, the results show a decrease in performance, with the average accuracy dropping from 99% to 50.6%. Also similar to our findings with hematopoiesis, changing the model’s architecture (LSTM vs. GRU, hyperparameter changes) has a limited impact on accuracy, which remains in the range of approximately 49% to 51%. This again highlights the importance of considering clonal structure in accurately predicting reprogramming outcomes (Supplementary Figure 5b, Supplementary Table 3).

### Deep Lineage can identify key genes in cellular reprogramming and generate realistic in silico perturbation experiments

Detecting and targeting key genes involved in cell reprogramming is useful for advancing our understanding of the system and potential therapeutic applications (21). Deep Lineage uses its learned latent representation to predict cellular fate and gene expression profiles and SHAP analysis to identify key genes within the reprogramming process. We tested this ability of Deep Lineage by using both fibroblast reprogramming and mouse hematopoiesis data (18,19) and performed a series of in silico perturbations to manipulate the fate of these systems. Using SHAP, we predicted key genes for the reprogrammed and failed trajectories across various time points. Known essential marker genes of reprogramming (19,22–24) (Apoa1, Cryab, Col1a1, and Sox11) are in this list. We also performed Gene Set Enrichment Analysis (GSEA) on the top 200 gene candidates demonstrating the ability of Deep Lineage to detect reprogramming-relevant biological pathways such as system and tissue development, proliferation signals, and cellular differentiation (Fig. 6c). Together these results suggest that Deep Lineage can be used to identify important reprogramming genes and processes.

**Fig. 5:**
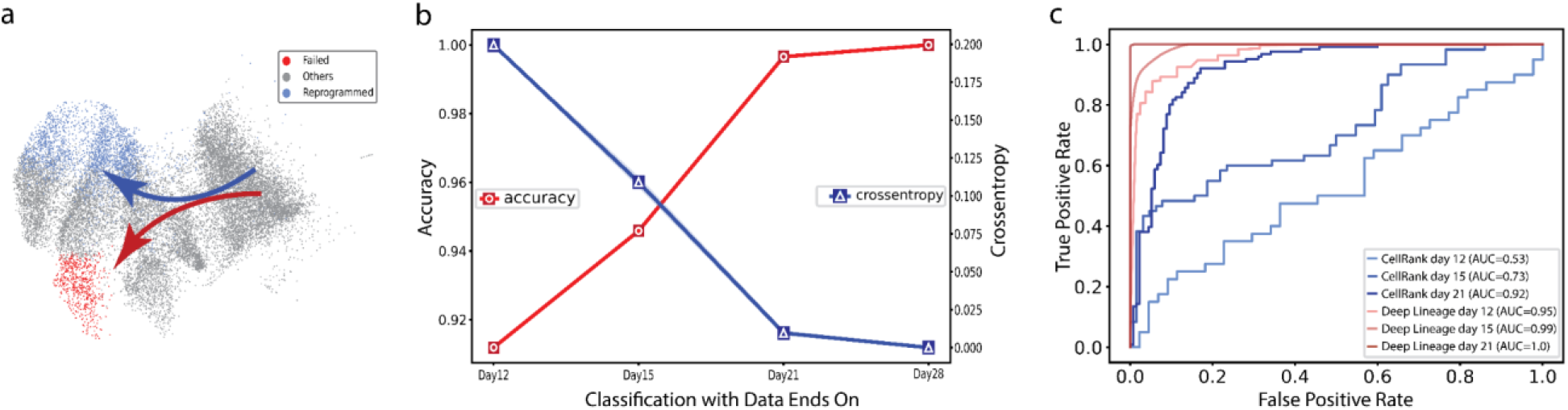
a, UMAP visualization illustrates progenitor bias toward either successful or failed reprogramming outcomes. b, The model’s accuracy in detecting successful or failed outcomes of progenitors is examined using gene expression data of cells up to days 12, 15, 21, and all days within a clone. c, Benchmarking comparison between Deep Lineage and CellRank in predicting fate outcome when using all time points up to a given time point (e.g. day 12) to infer time point “day 28”. Using ROC curves and AUC values, the graph shows the predictive power of both models in predicting fate bias. Deep Lineage consistently outperforms CellRank.

**Fig. 6:**
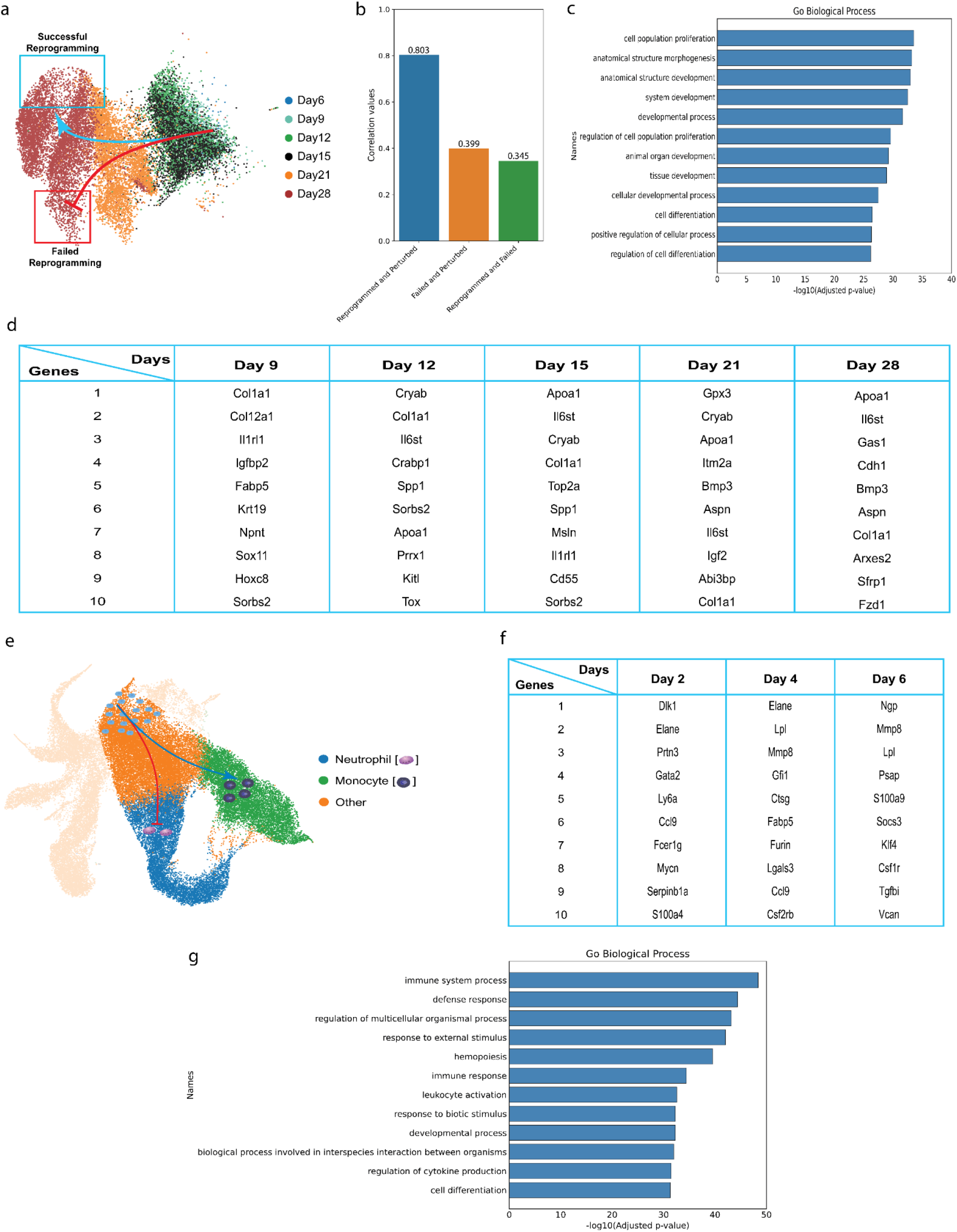
(a) UMAP representation showing the procedure for in silico perturbation of cells on day 15 within failed reprogrammed clones, aiming to switch them into successfully reprogrammed states. b, Correlation values between in silico perturbed data and actual successfully and unsuccessfully reprogrammed cells on Day 28, showing the power of Deep Lineage in performing in silico experiments. c, Gene set enrichment analysis reveals the biological processes associated with the top 200 genes identified by the SHAP method (40 genes selected from each time point) d, Identification of the top 10 gene candidates using SHAP analysis at each time point during fibroblast cell reprogramming. e, UMAP visualization depicting the in silico perturbation experiment targeting progenitor cells (Dark orange) in clones with a Neutrophil fate (blue), resulting in a transformation to Monocytes (green). f, The top 10 gene candidates for hematopoiesis detected by SHAP analysis across all time points. g, Gene set enrichment analysis of the top 200 genes identified by SHAP analysis.

Our goal is to alter cellular fate and reprogram cells in silico, steering them away from trajectories heading towards undesirable “dead-end” outcomes (Fig. 6a). We focused on the top 10 genes identified by the SHAP method in the failed trajectory path (Fig. 6d). After conducting a series of in silico perturbation experiments, which included over-expression and knock-out of these genes on day 15 of failed outcome clones, we applied Deep Lineage to generate gene expression profiles for days 21 and 28 (Fig. 6a). Subsequently, we input the generated perturbed time series back into Deep Lineage to evaluate the perturbations’ effectiveness in redirecting the fate of the initially failed clones. After the perturbation, the classifier shows that the trajectory changed, with the fate of the failed clones changing to successfully reprogrammed, demonstrating the ability of the model to predict a way to lead to more successful cellular reprogramming. To further evaluate these predictions, we conducted a correlation analysis comparing the average gene expression of in silico reprogrammed cells on day 28 to both real reprogrammed cells and real failed cells. Our results show a significantly higher similarity between the generated in silico successful reprogrammed cells and real successful reprogramming cells (Pearson correlation = 80.3%) compared to the real failed cells (39.9%) (Fig. 6b). This additional analysis further supports Deep Lineage’s ability to generate realistic gene expression for cells undergoing in silico perturbation.

We also applied Deep Lineage to hematopoiesis data to identify important genes and simulate the fate switch from neutrophils to monocytes (Fig. 6e). This analysis identified important genes, including some known genes (e.g. Dlk1, Gata2, Mmp8, Elane, and Lpl) important in the switch from neutrophils to monocytes (Fig. 6f) (25–29). Using the trained model and performing GSEA on the top 200 identified genes reveals significant enrichment of pathways associated with immune system development, differentiation, and hematopoiesis (Fig. 6g), indicating the model’s ability to capture biological pathways relevant to hematopoiesis. By performing in silico knockouts of the top genes of progeny cells on day 2 of clones predestined to become neutrophils, we were successfully able to alter the predicted final fate of the clone, pushing it toward monocytes by day 6. These results again demonstrate that Deep Lineage can accurately simulate and predict gene expression during in silico perturbation.

## Discussion

The analysis of cellular trajectories during differentiation is useful for understanding developmental processes and supporting the development of stem cell therapies. By investigating the routes cells take during these processes, we gain insights into cellular decision-making mechanisms and the timing of critical events (30,31). Computational methods for mapping cellular trajectories, such as pseudotime inference, optimal transport, and latent embedding models only consider gene expression information, even if clonal information is available (5,32–34). To consider both gene expression and clonal information in trajectory analysis, we developed Deep Lineage, a novel method combining autoencoder-based representation learning and Long Short-Term Memory (LSTM) or Gated Recurrent Units (GRUs) models for sequential analysis that capture how cells are related to each other within a clonal lineage. Deep Lineage can accurately predict gene expression and early cell fate bias as well as key genes for cellular reprogramming. It also can generate accurate in silico predictions of the results of perturbation (e.g. knockout and overexpression) experiments.

Deep Lineage implements three neural network architectures to model sequential clonal information: bidirectional LSTM, LSTM and GRU. In our results, bidirectional LSTM networks exhibit the highest accuracy in both predicting gene expression at different days and identifying early fate bias but also have the highest computational cost. LSTM provides an intermediate option, providing close to bi-LSTM accuracy with less cost, and GRU has the least computational cost, but sacrifices a small amount of accuracy compared to LSTM. Supporting these three architectures enables researchers to balance accuracy and computational efficiency, facilitating wider adoption in scenarios with limited computational resources.

Deep Lineage identifies key genes that may be useful to help reprogram cells. However, we cannot prove that our model is causal, as there are not enough perturbation experiments available to train or test. Thus model predictions of key genes is hypothesis hypothesis-generating and can be used with additional information, such as expert knowledge, to select genes more likely to be causally implicated in reprogramming processes.

Current single-cell lineage tracing experiments implement one-step or cumulative barcoding. In one-step, a single DNA barcode is introduced early and traced over time in a developing system, which in cumulative barcoding, new barcodes are generated as the system develops (13). We found the inclusion of additional barcoding information in the reprogramming data (cumulative barcoding) contributes to higher prediction accuracies compared to the hematopoiesis data (one-step) (0.93 vs. 0.81). This suggests that cumulative barcoding captures a more comprehensive picture of cellular lineage, leading to enhanced model performance in single-cell trajectory analysis.

We applied Deep Lineage to explore the dynamics of regenerative and developmental processes. However, it should be possible to use it to study other types of cellular plasticity, such as tumour development. Lineage tracing in tumours can be accomplished by inferring clonal relationships between cells based on somatic mutation profiles, as new mutations are generated with every cell division, and this process is typically amplified in cancer (35). Additionally, Deep Lineage’s in silico perturbation simulation capability can be used to explore novel treatment strategies and drug interventions. Future work can also expand Deep Lineage by integrating additional data modalities such as epigenomic data.

We envision Deep Lineage as a valuable tool for investigating the complex trajectories in regeneration and various developmental processes where the system experiences diverse dynamics.

## Methods

### Autoencoder for latent embedding of scRNA-seq data

To achieve the reduced dimension of the data before training the LSTM model, we employed an autoencoder, an unsupervised neural network algorithm (36). The autoencoder consists of two components: an encoder, which compresses the high-dimensional scRNA-seq data into a lower-dimensional latent space, and a decoder, which reconstructs the data back to the original input space. The objective of the autoencoder is to minimize the reconstruction error, measured using the mean squared error (MSE) loss function, expressed as:

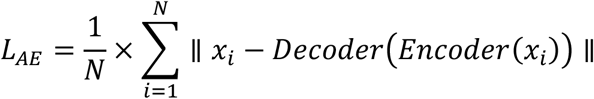

Where N is the number of training samples, *x*_*i*_ is the i-th input data sample which is the gene expression of a cell. *Encoder()* and *Decoder()* represent the encoder and the decoder parts of the autoencoder. By minimizing the above loss, the autoencoder learns a lower-dimensional representation of the data while preserving important biological features, facilitating the analysis of time series models (LSTMs, GRU).

### RNN models for gene expression and early cell fate prediction

For the regression task of predicting gene expression for cells within a clone, we harnessed the power of RNNs, mainly LSTMs and GRU. Unlike traditional feedforward neural networks, LSTM networks have loops and gates that enable information to persist and flow through time steps, making them well-suited for tasks involving time series and sequential data analysis. The distinguishing feature of LSTM is its memory block, which maintains a hidden state and a block state. The block state acts as a long-term memory, enabling the network to keep and update relevant information over long time ranges. The hidden state carries short-term memory and interacts with the block state through various gates. The LSTM block is composed of three main gates that control the flow of information: the input gate (i), the forget gate (f), and the output gate (o) (37). The input gate decides how much of the new input should be added to the block state. The forget gate controls what information from the previous block state should be discarded, allowing the network to selectively remember or forget information. The output gate determines what part of the block state should be exposed as the hidden state. The equations governing the LSTM block operations are as follows:

Input gate *i*_*t*_:

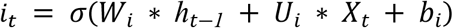

Forget gate *f*_*t*_:

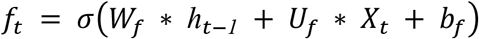

Output gate *o*_*t*_:

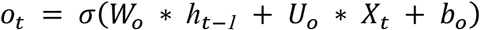

Block state *c*_*t*_:

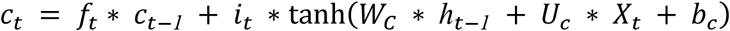

Hidden state *h*_*t*_:

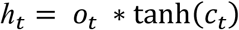

Here, *X*_*t*_ represents the input at the current time step, *h*_*t−1*_ denotes the hidden state from the previous time step, and σ is the sigmoid activation function. The weights and biases, represented by *W*_*i*_, *W*_*f*_, *W*_*o*_, *W*_*C*_, *U*_*i*_, *U*_*f*_, *U*_*o*_, and *U*_*c*_, are learnable parameters that the LSTM network optimizes during training. To obtain a compact representation of the gene expression data, we used the latent layer of an autoencoder as input to the LSTM model.

GRU is another variant of the RNN models, designed to address some of the limitations of traditional RNNs, such as the vanishing gradient problem that can interfere with the learning process. Similar to LSTM, GRU is particularly effective in processing sequential data and capturing dependencies over long sequences. Each GRU block also contains a hidden state, but instead of having separate blocks and hidden states like LSTM, it uses an update gate (z) and a reset gate (r) to control the flow of information (37). These gates enable the GRU to selectively update and reset the hidden state, facilitating the retention of relevant information and reducing the number of parameters compared to LSTM. In GRUs the reset gate controls how much of the previous hidden state should be forgotten, allowing the network to decide what information to retain from the past. The update gate determines how much of the candidate’s hidden state *h’*_*t*_ should be blended with the previous hidden state *h*_*t−1*_ to generate the new hidden state *h*_*t*_. This gate mechanism enables the GRU to regulate the information flow effectively. The operations of a GRU block can be summarized as follows:

Reset gate *r*_*t*_:

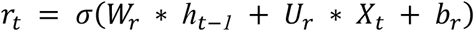

Update gate *z*_*t*_:

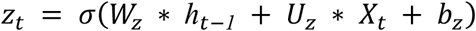

Candidate hidden state *h’*_*t*_:

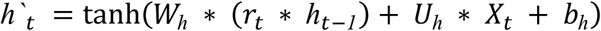

Hidden state *h*_*t*_:

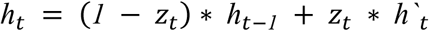

Here, *X*_*t*_ represents the input at the current time step, *h*_*t−1*_ denotes the hidden state from the previous time step, and σ is the sigmoid activation function. The weights and biases, are represented by *W*_*r*_, *W*_*z*_, *W*_*h*_, *U*_*r*_, *U*_*z*_, *U*_*h*_, *b*_*r*_, *b*_*z*_, and *b*_*h*_ are learnable parameters that the GRU network optimizes during training. GRU is computationally more efficient than LSTM due to its simplified architecture, making it particularly suitable for handling extensive datasets with limited computational resources or situations where training is scarce.

The equations mentioned earlier can be used for both regression and classification tasks. In classification tasks, the model’s output is then connected to a fully connected layer that handles the classification. Employing softmax activation in this layer allows the model to derive class probabilities.

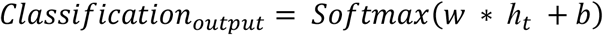

### Data preprocessing

Given the selection of a preprocessing method depends on the specific goal of the analysis, the performance of a supervised method can serve as a useful metric. Therefore, in each task (prediction of cell fate and gene expression), we compare the performance of different preprocessing steps before training the models and choose the one that leads to higher accuracy (lower reconstruction error for AE and lower loss for RNN) for that data. Our evaluation involves a multitude of combinations that incorporate different count normalization, log transformation, several scaling techniques, different numbers of highly variable genes, and different dimensionality reductions. For both reprogramming and hematopoiesis data, we realize that log transformation without count normalization achieves the highest accuracy and as expected using only raw counts or just scaling without log transformation results in poor performance which is consistent with the recently published benchmark paper (38) (Supplementary Figure 2, Supplementary Table 1). Following this, we select the highly variable genes and employ an autoencoder for dimensionality reduction (the architectures can be found in Supplementary Table 1).

### Modelling choices and training

After completing the preprocessing step, the gene expression data is partitioned into three subsets: 80% for training, 10% for validation, and 10% for testing. To optimize model performance, we conduct an extensive hyperparameter search using various model types (LSTM, GRU, bidirectional LSTM), model architecture (number of layers, nodes in each layer), and activation function (Tanh, ReLU, Leaky ReLU with alpha=0.3) (the detailed results are provided in Supplementary Table 2). Subsequently, we select specific sets of parameters that demonstrate superior accuracy for both the autoencoder reconstruction and the LSTMs during regression and classification tasks, tailored to each dataset. For the autoencoder, we employ the mean square error as the loss function, to ensure effective learning of the latent representation. After training the autoencoder, the resulting latent embedding is used to train the RNN models. In the regression task, where we aim to predict gene expression levels of cells across different developmental stages, we employ the mean square error as the loss function. For predicting early cell fate bias (classification task), we adopt the categorical cross-entropy loss function. Throughout the training process of both the autoencoder and the RNN models, we implement early stopping, which enables us to halt the training once the models demonstrate optimal performance on the validation set, preventing overfitting and supporting generalization to unseen data. In each iteration, we randomly select a training sample and update the network’s weights using SGD (Stochastic Gradient Descent) and Adam optimization algorithms with a default learning rate of 0.001. Also, RNN training can be slow due to the sequential nature of computations. To tackle this challenge, we use the CuDNN LSTM technique, which uses GPU-accelerated optimization to enhance the training efficiency of Long Short-Term Memory (LSTM) networks. By using the parallel processing capabilities and specialized algorithms offered by CuDNN LSTM, we achieve a significant acceleration in model training speed. In addressing instances of missing data within time series, a common approach involves the use of masking techniques. When a data point at a specific timestep is not available, the model skips that particular timestep and proceeds to the subsequent one. By using masking in both classification and regression tasks, we ensure that the time series model handles missing data consistently and effectively. Regarding the model selection, we observe GRU, and LSTMs achieve high accuracies with LSTMs showing slightly superior performance (Supplementary Figure 2, Supplementary Table 1). However, if computational resources are limited, we recommend using GRU over LSTMs, as GRU requires less computing power.

### Comparison with existing methods

To establish a benchmark for evaluating the performance of Deep Lineage in early cell fate bias and predicting gene expression of cell progenies, we selected existing relevant methods to compare to: Cospar (9), Super OT (10), WOT (8), and GAN-Based OT (10). We employ the same task as performed by Super OT (10), using the accuracy metrics reported in their study, thus enabling a direct comparison of our model’s performance against these established benchmarks.

In the context of early cell fate bias in the reprogramming dataset, we conduct a comparative analysis between Deep Lineage and Cellrank (20). The Cellrank framework integrates similarity-based trajectory inference with RNA velocity to uncover directed probabilistic state-change trajectories. When applying Cellrank to the reprogramming data, we followed the recommended preprocessing steps, including filtering genes based on a minimum expression threshold across ten cells and at least 20 counts in both spliced and unspliced layers. Following this, normalization by total counts per cell was performed, along with log transformation, while selecting the top 2,000 highly variable genes. Subsequently, a PCA representation was computed using the top 30 principal components (PCs) to construct a K-nearest neighbor (KNN) graph, with the parameter K set to 30.

### Hardware used

Two computational setups are used for data processing, model training, and plotting. A distributed computing system was used extensively for data processing and model training. Nvidia A100 (40GB) and Nvidia P100 (16GB) GPUs were used on this computing server. Intel Xeon E5-2680v4 2.4 GHz (Broadwell) and AMD EPYC 7542 2.9 GHz CPUs were used for other tasks. The maximum memory used on these servers was 64GB.

## Data availability

The data used in this study are openly accessible and can be obtained from their respective public repositories. The hematopoiesis Weinreb et al. data is available for download from the Gene Expression Omnibus (GEO) database under accession number GSE140802. The reprogramming dataset can be accessed from the GEO database with accession number GSE99915. Additionally, a preprocessed version of the data can be obtained from: https://cospar.readthedocs.io/en/latest/.

## Code availability

Code supporting this study is available on:…. We used Python 3.9.13, tensorflow-gpu 2.10.0, cudatoolkit 11.3.1, cudnn 8.4.1.50, matplotlib 3.5.1, numpy 1.22.4, scanpy 1.8.2, scikit-learn 1.1.1, seaborn 0.12.2, scipy 1.10.1, keras 2.10.0, cospar 0.2.1, pip 22.1.2, pandas 1.4.3, conda 22.9.0.

## Notes

### Competing Interest Statement

The authors have declared no competing interest.

